# Predictive modeling of single-cell DNA methylome data enhances integration with transcriptome data

**DOI:** 10.1101/2020.06.05.137000

**Authors:** Yasin Uzun, Hao Wu, Kai Tan

## Abstract

Despite rapid advances in single-cell DNA methylation profiling methods, computational tools for data analysis are lagging far behind. A number of tasks, including cell type calling and integration with transcriptome data, requires the construction of a robust gene activity matrix as the prerequisite but challenging task. The advent of multi-omics data enables measurement of both DNA methylation and gene expression for the same single cells. Although such data is rather sparse, they are sufficient to train supervised models that capture the complex relationship between DNA methylation and gene expression and predict gene activities at single-cell level. Here, we present MAPLE (Methylome Association by Predictive Linkage to Expression), a computational framework that learns the association between DNA methylation and expression using both gene- and cell-dependent statistical features. Using multiple datasets generated with different experimental protocols, we show that using predicted gene activity values significantly improves several analysis tasks, including clustering, cell type identification and integration with transcriptome data. With the rapid accumulation of single-cell epigenomics data, MAPLE provides a general framework for integrating such data with transcriptome data.

## Introduction

Recent advances in single-cell genomics provide the opportunity to capture a variety of epigenomic signatures at single-cell resolution including histone modification (Grosselin et al. 2019), DNA methylation (Smallwood et al. 2014; Urich et al. 2015; Luo et al. 2017, 2018; Mulqueen et al. 2018), chromatin accessibility (Buenrostro et al. 2015; Chen et al. 2018) and long-range chromatin interaction (Nagano et al. 2013; Stevens et al. 2017). In particular, single-cell DNA methylome analysis can provide quantitative and high-resolution measurement of cell-type-specific epigenomic landscape in both development and disease, as the mammalian embryonic development is associated with dynamic changes in DNA methylation at *cis*-regulatory elements and genome-wide deregulation of DNA methylation is associated with many types of cancer (Greenberg and Bourc’his 2019).

Identifying genome-wide methylation signatures at single-cell resolution with an unbiased technique such as bisulfite sequencing (BS-seq) comes with unique challenges. Technically, the data is sparse and genomic coverage is rather limited (∼5% of the genome per cell on average), even for deeply sequenced samples with more than 5 million reads per cell (Luo et al. 2017). Biologically, the interpretation of methylome signal is context dependent. Whole genome DNA methylome data have shown that gene body methylation is positively correlated with gene expression in embryonic stem cells (Lister et al. 2009; Greenberg and Bourc’his 2019), whereas in other cell types such as post-mitotic neurons, genic methylation is negatively correlated with gene expression (Lister et al. 2013; Lee et al. 2019). These observations suggest that the gene regulatory roles of DNA methylation are both genomic feature and cell-type-specific. Single-cell multi-omics studies suggest significant correlation (both positive and negative) between expression and gene body methylation for only a limited number of genes (Hu et al. 2016; Angermueller, Clark, et al. 2016). As a repressive marker, mean promoter methylation is significantly negatively correlated with gene expression only in a fraction of promoters (Angermueller, Clark, et al. 2016; Clark et al. 2018; Argelaguet et al. 2019) in individual cells. As a result, there is not a straightforward approach for inferring gene activity levels using single-cell DNA methylation datasets.

The lack of well defined association between DNA methylation and gene expression poses obstacles for the analysis and integration of this type of epigenomic data. Identifying cell types is relatively straightforward in scRNA-Seq data using clustering techniques and marker genes. However, the same approach does not work well for single-cell methylation data due to the lack of clear association between methylation and gene expression. As a result, it is difficult to use marker genes to assign cell types in this case. In terms of data integration, multiple computational methods have been developed for integrating different types of single-cell data (Stuart et al. 2019; Welch et al. 2019; Korsunsky et al. 2019). However, since these methods are dependent on reliable gene activity scores as the input, robust estimate of methylation-based gene activity score is necessary and remains a bottleneck for accurate integration of DNA methylation data with other types of single-cell omics data, which in turn can pave the way for comparative analysis of gene regulation mechanisms across cell populations in a complex tissue.

Here, we describe Methylome Association by Predictive Linkage to Expression (MAPLE), a supervised learning framework for predicting gene activity score based on single-cell DNA methylation data. We develop both gene-dependent and cell-dependent statistical features as the input to an ensemble learning framework. To train the statistical predictor and to evaluate the accuracy of the predictions, we take advantage of datasets generated using true multi-omics protocols (i.e. joint profiling of transcriptome and DNA methylome of the same physical cells). It has already been demonstrated that several hundreds of cells can be sequenced for gene expression and DNA methylation in the same cell (Angermueller, Clark, et al. 2016; Clark et al. 2018). Although the coverage of such multi-omics data is rather shallow, we demonstrate that they are sufficient for training robust predictors. Finally, we show that the predicted gene activity score significantly enhances cell type identification and integration of single-cell methylation data.

## Results

### Overview of the method

We hypothesize that the common patterns of promoter methylation-gene expression association for groups of genes can be modeled by using a supervised learning model. We developed two classes of statistical features as input to the supervised predictor. Promoters overlapping with CpG islands or shores have distinctive response to methylation (Deaton and Bird 2011; Jones 2012; Weber et al. 2007; Greenberg and Bourc’his 2019) and overall promoter CpG frequency has been used as a feature for predicting gene expression using bulk DNA methylation data (Kapourani and Sanguinetti 2016). In MAPLE, we used CpG frequency at higher resolution, as a vector of CpG frequencies of multiple genomic bins tiled across the promoter. This step constitutes the gene-dependent but cell-independent feature set to be used in the learning model. For the gene- and cell-dependent feature set, we computed the methylation level of each promoter bin for all cells and genes.

The sparsity of single-cell bisulfite sequencing data poses a significant challenge for machine learning approaches. Many promoter regions have a limited number of CpG sites covered in each single cell and dividing the promoter region into multiple bins further exacerbates the sparsity problem, resulting in either bins with no overlapping calls at all, or insufficient calls to make a reliable estimation of the methylation level of the bins. To alleviate this problem, we resorted to the concept of ‘meta-cell’, essentially borrowing information from neighboring single cells (Wagner, Yan, and Yanai 2017; van Dijk et al. 2018; Gong et al. 2018; Zhu et al. 2019). Each meta-cell represents a group of individual cells that are in a similar state with a specific cell in the data (Fig. 1a, Methods). Although this approach results in a slight loss of data resolution, it provides a reliable estimation of CpG level for the vast majority of meta-cells, even with small neighborhood sizes (Supplemental Fig. 1). Moreover, each single cell has its own unique neighborhood, thus no two meta-cells are expected to be identical and the single-cell nature of the data is preserved.

**Figure 1.**
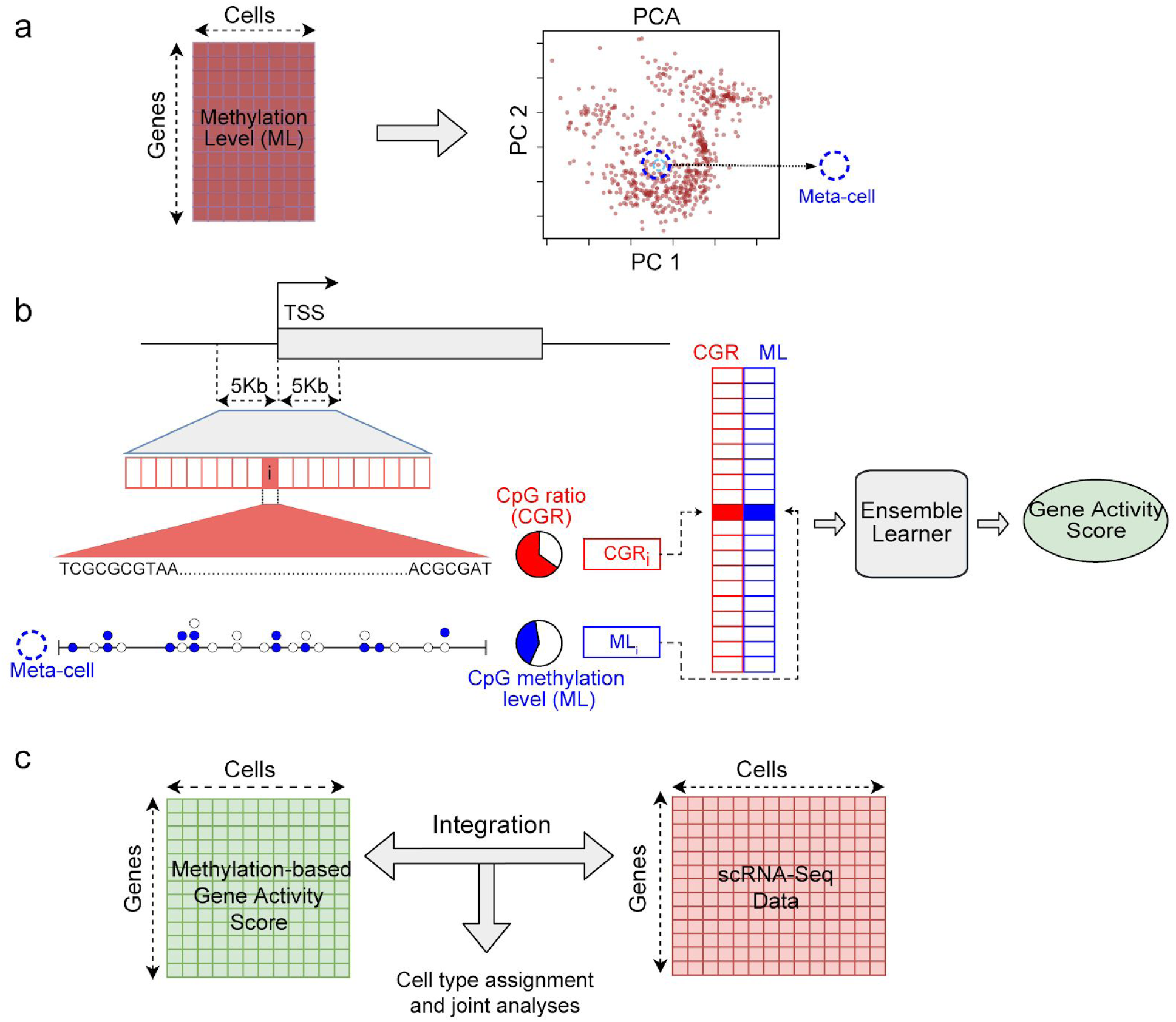
Schematic overview of the computational framework. **a)** Generation of meta-cells from single-cell DNA methylation data. The gene-by-cell DNA methylation matrix is used for principle component analysis (PCA) to reduce the dimensionality. Each point in the PCA plot is a single cell. A meta-cell is the set of *k* nearest cells to an individual cell in the PCA space. **b)** Prediction of gene activity level by combining DNA methylation and sequence information in promoter. TSS, Transcription Start Site. ML, Methylation Level. Each horizontal box represents a genomic bin. Each circle represents a CpG site and filled circle represents methylated cytosine and empty circle represents unmethylated cytosine in a meta-cell. CpG ratio is defined as the percentage of CpG dinucleotides in a genomic bin. CpG methylation level is defined as the ratio of the number of methylated CpG calls to all CpG calls. **c)** Integration of single-cell methylation and single-cell RNA-Seq data based on predicted gene activity scores using single-cell DNA methylation data.

Once the meta-cells are identified, the CpG methylation level for each meta-cell is computed for 500bp bins across the promoter region, resulting in 20 features for each gene and each meta-cell. Together with the CpG frequencies of these bins, they constitute the two classes of features for each gene-meta-cell pair (Fig 1b, Methods). These features are used as the input to the supervised learning model to infer the gene activity level in the meta-cell. For downstream analysis, a DNA methylation-based gene activity matrix is constructed and a desired method can be used to integrate the methylome data with gene expression data, for reliable assignment of cell types in the data and other joint multi-omics analyses (Fig. 1c).

Many supervised learning methods can be used to predict the gene activity level of the cells. To achieve a consistent performance independent of training and test data, we applied an ensemble approach by combining multiple predictors (Mendes-Moreira, Soares, and Jorge 2012). We selected one commonly used, representative predictor from three broad categories: artificial neural networks, regression-based models and decision tree based models. Convolutional Neural Networks (CNNs) is a class of deep-learning neural networks, which have gained popularity in image processing and bioinformatics (Angermueller, Pärnamaa, et al. 2016; Alipanahi et al. 2015; Angermueller et al. 2017). Our CNN architecture resembles the method described in (Singh et al. 2016) for predicting gene expression using bulk histone mark ChIP-Seq data. However, instead of multiple histone marks, we used two classes of features, namely, CpG dinucleotide frequencies and CpG methylation levels for genomic bins surrounding the TSS (Fig. 1b, Methods). Elastic Net (EN) is a regularized regression method that combines the least absolute shrinkage and selection operator (Lasso) and ridge regression models. Random Forest (RF) consists of multiple decision trees, each of which is trained with random subsamplings from the training data and the result is obtained by combining the outputs of all decision trees in a democratized manner. As the baseline method, we computed the mean promoter de-methylation level (MPD) (ratio of unmethylated CpGs to all CpG calls in the promoter) as a predictor of gene activity level.

### Supervised learning improves accuracy of gene activity prediction

We benchmarked the performance of MAPLE using four published single-cell multi-omics datasets generated using two different protocols. The datasets of Angermueller et al. (Angermueller, Clark, et al. 2016) and Hernando-Herraez et al. (Hernando-Herraez et al. 2019) were generated using scM&T-Seq (single-cell genome-wide Methylome and Transcriptome sequencing), while datasets of Clark et al. (Clark et al. 2018) and Argelaguet et al. (Argelaguet et al. 2019) were generated using scNMT-Seq (single-cell Nucleosome, Methylation and Transcription sequencing). For all datasets, the transcriptome and DNA methylome were jointly profiled for the same single cells.

There are different ways for combining component predictors to build an ensemble predictor, such as unweighted average, weighted average and stacking. We evaluated all three approaches and found that they have similar performance across all combinations of training and test datasets (Supplemental Fig. 2). We therefore chose the unweighted average approach due to its simplicity. To evaluate the overall performance of the ensemble approach, we performed external cross validation, i.e. training a predictor with one dataset and predicting the gene activity levels using the remaining three datasets. We evaluated the prediction accuracy using two metrics, Spearman correlation and median squared error. We chose Spearman correlation because it can capture both linear and nonlinear relationships in the data. Across the twelve training-test datasets, MAPLE achieved an average global Spearman correlation of 0.62 (across all genes and cells). In comparison, the baseline predictor using MPD as the feature gave an average global Spearman correlation of 0.5 (Fig. 2a). The Spearman correlation across genes is also higher for MAPLE compared to the baseline predictor (Fig. 2b, Supplemental Fig. 3). We observed the same trend when using a median squared error as an alternative metric (Supplemental Fig. 4, 5). Taken together, these results demonstrate that a supervised predictor using both CpG frequency and methylation level can substantially enhance the accuracy of predicting gene expression activity at single-cell level.

**Figure 2.**
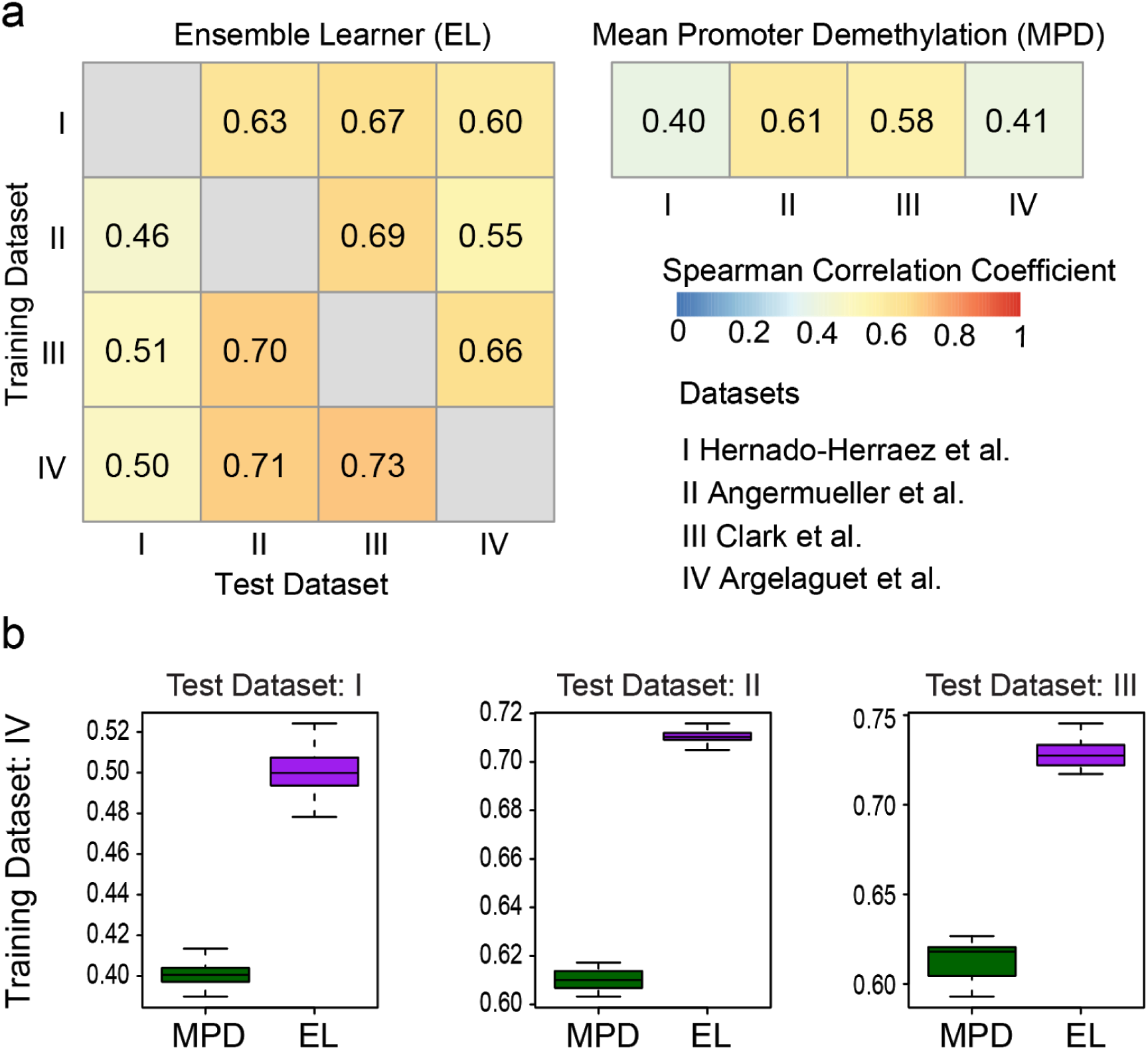
Prediction accuracy of gene expression using DNA methylation data and ensemble learning. **a)** Heatmap showing global Spearman correlation coefficients between observed gene expression and predicted gene activity for all genes across all cells in a dataset. Rows represent training datasets and columns represent test datasets. Row and column Roman numbers correspond to the datasets shown. **b)** Distribution of Spearman correlation coefficients across genes using the dataset of Clark et al. as the training set. Each data point represents one cell.

**Figure 3.**
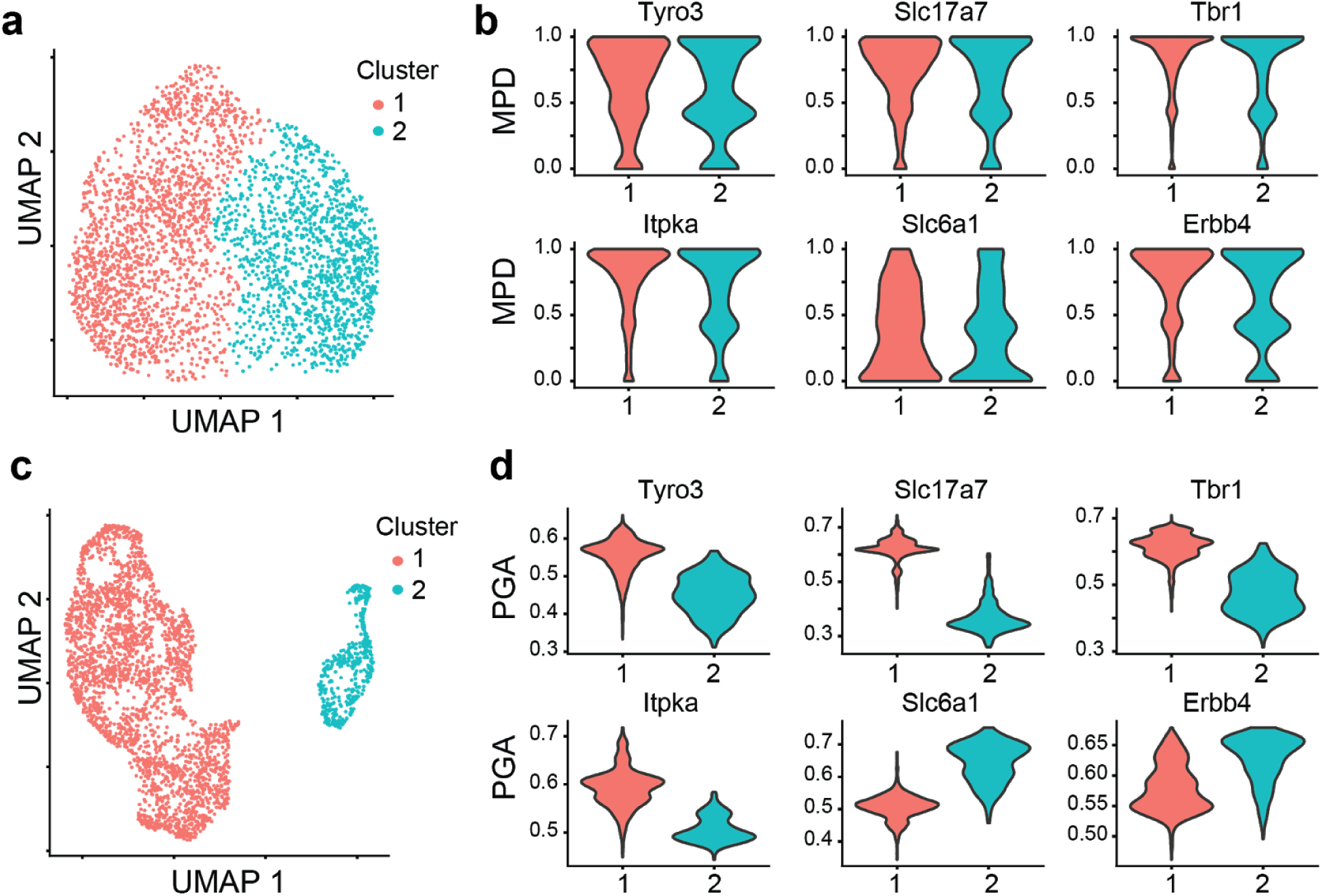
Predicted gene activity using methylome data improves cell type identification. **a)** UMAP of clustering result generated using mean promoter de-methylation (MPD) as the input. **b)** Violin plot of MPD values for marker genes for excitatory (*Tyro3, Slc17a7, Tbr1, Itpka*) and inhibitory (*Slc6a1, Erbb4*) neurons. **c)** Same as panel a) but using predicted gene activity (PGA) as the input. **d)** Same as panel b) but using predicted gene activity as the input.

**Figure 4.**
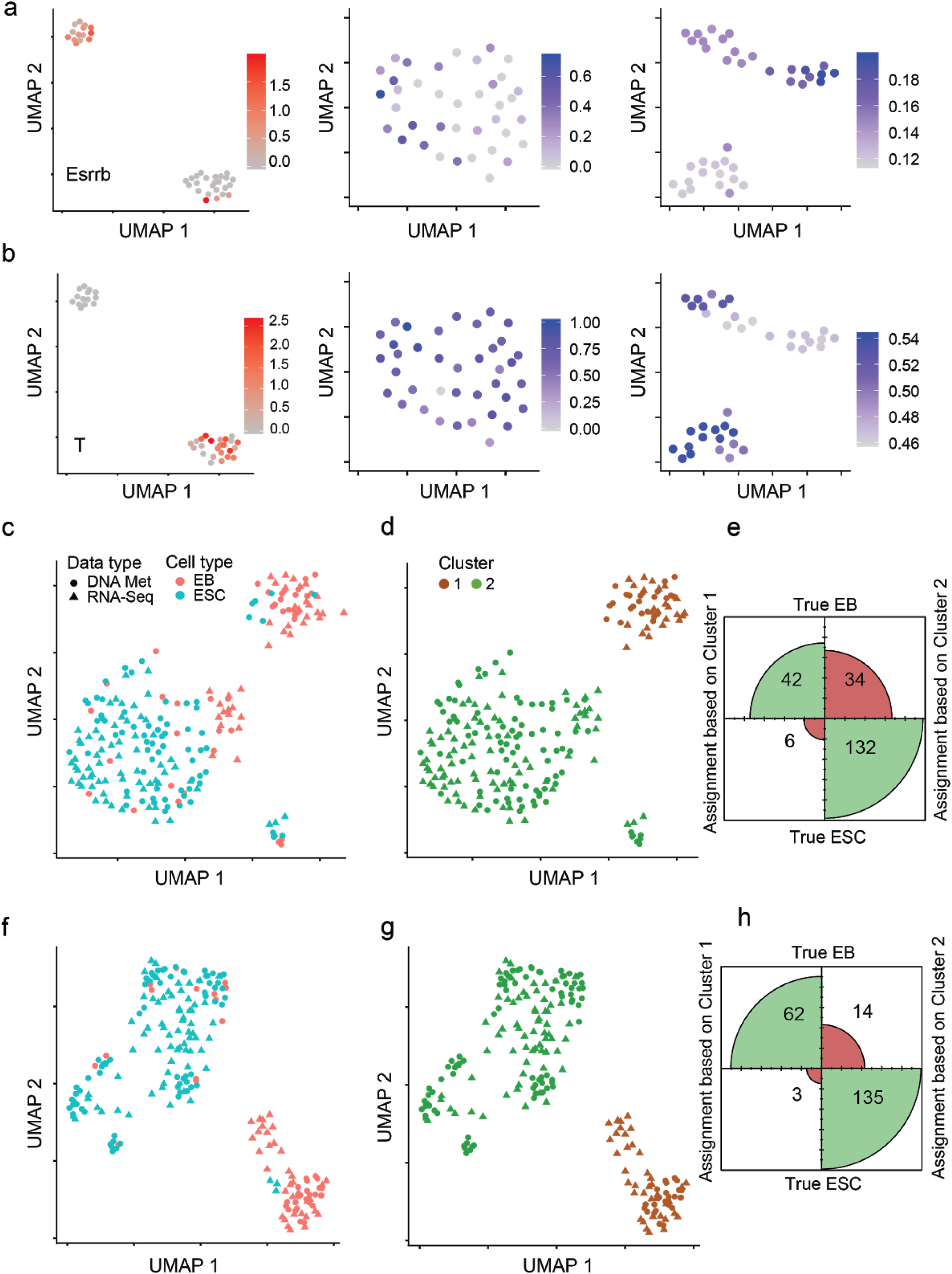
Predictive modeling improves integration with transcriptome data of cell lines. **a)** Cell heterogeneity based on transcriptome and DNA methylome data. Left panel, UMAP using RNA-Seq data as the input. Color scale represents the expression level of *Esrrb* for EBs. Middle panel, UMAP using mean promoter de-methylation (MPD) as the input. Color scale represents the MPD level of the *Esrrb gene*. Right panel, UMAP using predicted gene activity based on DNA methylation data as the input. Color scale represents the predicted gene activity levels of *Esrrb*. **b)** Same as panel a) but for the *T* gene. **c)** UMAP based on integrated RNA-Seq and DNA methylation data. Mean promoter de-methylation (MPD) was used as the input for data integration using Seurat. EB, embryoid body. ESC, embryonic stem cell. **d)** Density clustering of the data shown in the UMAP in panel c). **e)** Confusion matrix plot based on the clustering result shown in panel d), illustrating the agreement between cell type assignment based on clustering and true cell type. Size of each quadrant is proportional to the number of cells classified. **f)** Same as panel d), but using predicted gene activity as the input. **g)** Same as panel d), but using predicted gene activity as the input. **h)** Same as panel e), but using predicted gene activity as the input.

**Figure 5.**
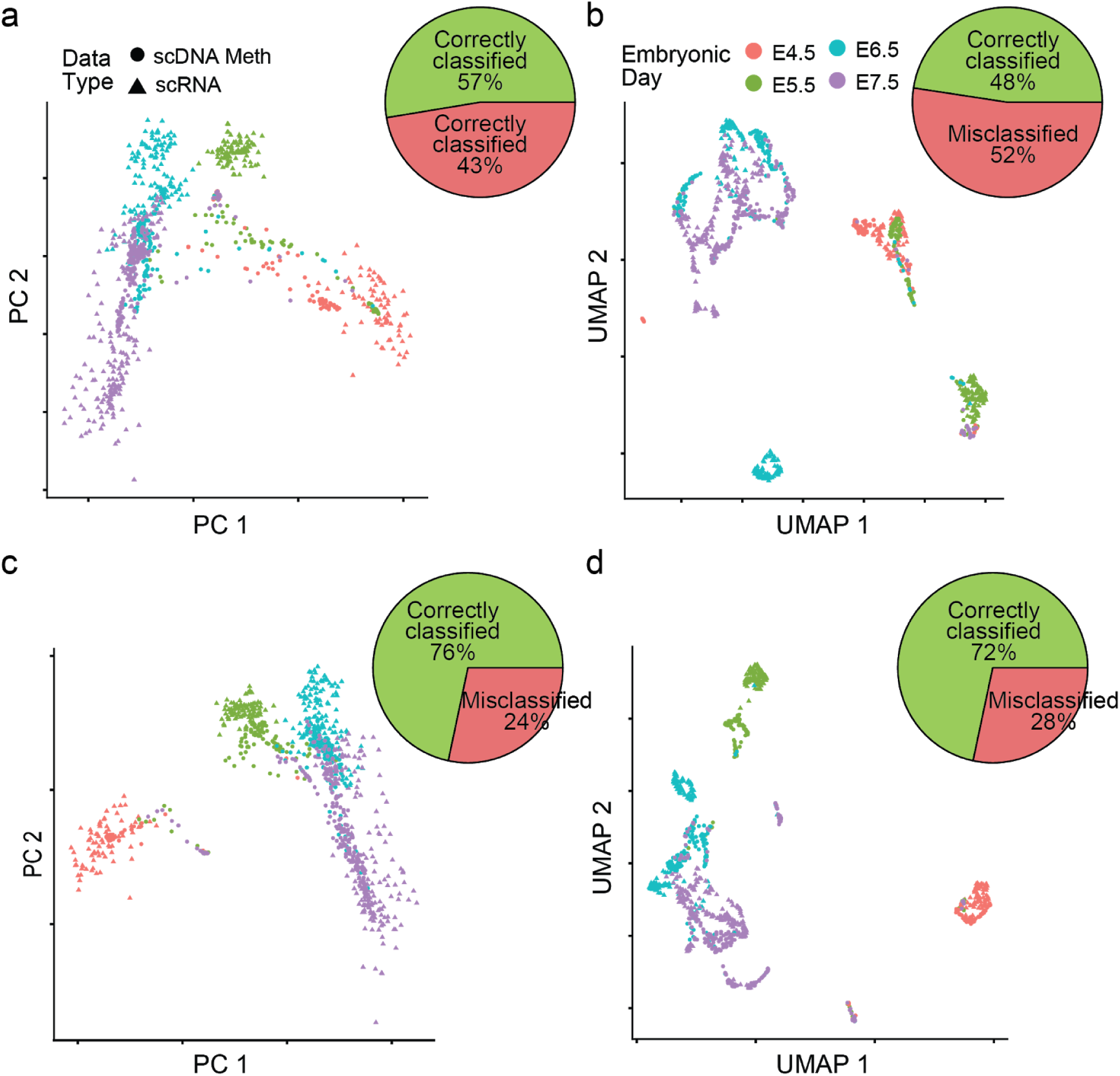
Predictive modeling improves integration with transcriptome data of primary tissues. **a)** Principal component analysis using integrated RNA-Seq and DNA methylation data. Mean promoter de-methylation (MPD) was used as the input for data integration using Seurat. Pie chart shows the percentage of correctly and mis-classified cells using scDNA-methylation data, based on k-nearest neighbor (kNN) classification on the scRNA-Seq cells. **b)** Same as panel a), but using UMAP instead of PCA as the dimensionality reduction method. **c)** Same as panel a), but using predicted gene activity as the input. **d)** Same as panel b), using predicted gene activity as the input.

### Predicted gene activity improves identification of cell types

Since one of the most important utilities of the single-cell data is to identify different cell types/states in a heterogeneous population, we next evaluated our trained predictor on a single-cell DNA methylation dataset containing 3,377 mouse neurons which were sequenced with the single-nucleus methylcytosine sequencing (snmC-seq) protocol (Luo et al. 2017). In this study, the authors identified different neuronal subtypes using gene body non-CpG (mCH) methylation, which is known to be inversely correlated with the expression level in adult neurons (Mo et al. 2015).

When we used the CpG (instead of non-CpG in order to be general) MPD value for dimensionality reduction, we observed no separation of cell types in the data (Fig. 3a) and the MPD values of the marker genes for the neuronal subtypes were indistinguishable across the cell populations (Fig. 3b). In contrast, when we clustered the cells using the gene activity levels predicted by MAPLE (trained on data from (Clark et al. 2018)), we identified two main clusters that were clearly separated (Fig. 3c). The larger cluster shows higher gene activity for excitatory neuron markers, such as *Tyro3, Slc17a7, Tbr1, Itpka* and the smaller cluster shows higher gene activity for inhibitory neuron markers, such as *Slc6a1, Erbb4* (Fig. 3d). The clustering result was highly consistent with that reported by the authors, with over 99% of the cells in the excitatory neuron cluster and 95% of the cells in the inhibitory cluster matching those reported by the original study, respectively.

To further evaluate the utility of predicted gene activity for cell type identification, we applied MAPLE to identify cell clusters in the embryoid body (EB) population in the multi-omics dataset by Clark et al. (Clark et al. 2018) using data from Angermueller et al. (Angermueller, Clark, et al. 2016) as the training set. In this study, the authors differentiated mouse embryonic stem cells to EB and reported that the differentiated population clusters into two main clusters, one is labeled as pluripotent due to high expression of pluripotency genes and the other represents differentiated cells with the opposite pattern. Using the MPD levels as the input for clustering did not recapitulate the heterogeneity observed using the transcriptome data. On the other hand, the same analysis using predicted gene activity by MAPLE as the input resulted in two clusters that have similar sizes to the clusters defined using transcriptome data alone. Moreover, *Esrrb*, a pluripotency marker gene showed high levels of activity in one cluster whereas transcription factor *T*, a differentiation marker, displayed the opposite pattern (Fig 4a, b), consistent with the pattern of heterogeneity based on the transcriptome data. In summary, these results demonstrate that use of a predictive modeling approach can significantly improve our ability to identify cell types and reveal heterogeneity in single-cell methylation data.

### Predicted gene activity enhances integration with scRNA-Seq data

We reasoned that accurate prediction of gene activity level can significantly enhance the integration of single-cell methylome data with transcriptome data. To test this hypothesis, we took advantage of published true multi-omics datasets in which DNA methylome and transcriptome were measured for the same single cells. Therefore, we know the ground truth about the matching of the two data types for a given cell. We first used the dataset by Clark et al. that is composed of differentiated embryoid body cells and undifferentiated embryonic stem cells (ESCs) (Clark et al. 2018). Using predicted gene activity levels by either MAPLE or the MPD method, we integrated the methylome and transcriptome data using Seurat (Stuart et al. 2019). (Stuart et al. 2019). We then clustered the cells using the co-embedded transcriptome and methylome data produced by Seurat. We computed the fraction of cells in the methylome data that were assigned to the correct cluster, using a k-nearest neighbor (kNN) classifier and cells from single cell RNA data. In contrast to MPD, clusters computed based on MAPLE-predicted gene activity showed higher homogeneity (i.e. greater fraction of the cells in each cluster belongs to the same type, Fig. 4c-h, Supplemental Fig. 6), suggesting improved accuracy in data integration.

We further evaluated the performance of data integration using a larger dataset on primary tissue. The dataset includes 850 cells from four different embryonic time points (Argelaguet et al. 2019) during mouse gastrulation. Each cell was sequenced with multiple modalities, including chromatin accessibility, DNA methylation and RNA expression. We integrated the single-cell methylome data with the expression data, as described above and computed the fraction of cells in the methylome data that were assigned to the correct cluster. Using gene activity predicted by MPD, neither linear (PCA) nor non-linear (UMAP) dimensionality reduction methods on the integrated data resulted in accurate matching of cells from the same developmental stage (Fig. 5a,b). In comparison, using MAPLE data as the input for integration, the resulting integrated data showed much higher fractions of matched cells based on the two data types for all four developmental stages (Fig. 5c,d, Supplemental Fig. 7). Although part of the populations from E6.5 and E7.5 has some overlap in the integrated data based on MAPLE input, this outcome is due to the biological nature of the data, as it is observed in all data modalities including gene expression (Argelaguet et al. 2019).

## Discussion

While recent advances in single-cell DNA methylome sequencing technologies improve our ability to study epigenetic heterogeneity, data analysis poses unique challenges, especially the issue of connecting the methylome and transcriptome data. Here, we addressed this challenge by developing a supervised learning approach to inferring gene activity based on DNA methylation data. The inferred gene activity score acts as an intermediate input for integrating the two types of data. A similar approach is commonly used to integrate single-cell ATAC-Seq data with transcriptome data where summed chromatin accessibility signal in the promoter and gene body is used as a proxy to gene activity and intermediate input for data integration. However, simply using summed DNA methylation signal in the promoter region does not work as we demonstrated using the mean promoter demethylation method, due to the more complex relationship between DNA methylation and gene expression. Instead, we showed that predictive modeling better captures the relationship between DNA methylation and gene expression. As a result, the predicted gene activity score helps improve cell type identification in the methylome data as well as integration with transcriptome data.

We used an ensemble learning approach to predict gene activity. Although we chose three representative predictors as components of the ensemble predictor, more predictors can be explored in the future. Similarly, additional ensemble combination rules can be explored to improve the accuracy.

In our framework, we used promoter methylation since it can be directly associated with genes. It has been reported that gene-distal transcriptional enhancers also have unique DNA methylation signature (Lister et al. 2009; Stadler et al. 2011; Hon et al. 2013). We anticipate that inclusion of enhancer DNA methylation signals can further improve the accuracy of the predictive model. To this end, a major challenge is associating distal enhancers with their target genes. This question has been addressed extensively for bulk data using both experimental and computational approaches (Jung et al. 2019; He et al. 2014; Javierre et al. 2016; Cao et al. 2017). However, due to unique challenges in single-cell data, additional research is warranted to address this important question.

## Methods

### Processing of public datasets

Datasets used in this study are listed in Supplemental Table 1. Briefly, we used four single-cell multi-omics datasets generated by two different experimental protocols, scM&T-Seq (Angermueller, Clark, et al. 2016) and scNMT-Seq (Clark et al. 2018). In addition, we used the snmC-Seq (Luo et al. 2017) dataset as the methylation-only data. We used Bismark methylation call files (cov files) when available from the authors, or converted the methylation calls into Bismark cov format (Krueger and Andrews 2011). For each dataset, we only used cells for which both scRNA-Seq and sc-Methylome data were available.

For the single-cell RNA-Seq data, we normalized the number of reads by the total number of reads per cell and obtained CPM (counts per million). CPM values were log transformed and were further normalized by the maximum log expression value in the dataset in order to fit all datasets into the same range and make training and test sets compatible.

### Computing meta-cells

In order to alleviate the problem of data sparsity, we combined DNA methylation data from neighboring cells into meta-cells as follows. We first counted the CpG methylation calls in the +/-5kbp region flanking the transcription start site (TSS) of the genes and computed the methylation level as the ratio of methylated CpGs to all CpG calls in those regions. For regions with no cytosine calls for a particular cell, we used the mean methylation level across all promoters of that cell. Next, we performed principal component analysis (PCA) on this methylation level matrix and computed the Euclidean distance between each cell pair. We used the top *d=10* principal components as the feature space to compute the distance (*d* is an adjustable parameter in our method), since the total variance explained after the 10th principle component was minimal (Supplemental Fig. 8). Based on the distance, we defined the local neighborhood of each cell as the (*k*-1) nearest cells, making a meta-cell size of *k* (*k* is an adjustable parameter in the method), including itself. Note that each meta-cell corresponds to an actual cell in the dataset, i.e. the number of meta-cells is the same as the number of single cells in the data. Since we observed that there is minimal improvement in terms of non-empty bins for *k* larger than 20 (Supplemental Fig. 9) for all datasets in this study, we used *k*=20.

### Calculation of DNA methylation rate

Transcription start sites (TSS) were defined based on the Gencode annotation (release vm23) for mouse genome (release GrCM38). Promoter was defined as the region spanning 5kbp up and downstream of the TSS.

To compute CpG methylation rate, we first divided each promoter into 20 500bp bins. For each meta-cell, the numbers of methylated and unmethylated CpG sites for each bin were counted. The methylation rate of each bin was calculated by dividing the total number of methylated cytosines to all cytosines in that bin, considering all the cytosine calls for all cells in the corresponding meta-cell.

### Feature Set

Two classes of features were computed, gene-dependent and cell-independent feature and cell- and gene-dependent feature. For each gene, we extracted the sequence information for the +/-5kbp region around the TSS. For each 500bp bin, we computed the CpG frequency, considering both strands, which resulted in 20 gene-dependent, cell-independent features. Cell- and gene-dependent features are CpG methylation rates of the 20 promoter bins of a gene in a given cell. They were calculated by using all methylation calls for all cells in the corresponding meta-cell.

### Training and testing the predictor

We randomly subsampled 100,000 cell-gene pairs from each training dataset in order to speed up the training process. Three supervised learners, convolutional neural network (CNN), elastic net (EN), and random forest (RF) were trained with each of the four multi-omics datasets. We evaluated the performance for different parameter settings for individual learners by using 5-fold internal cross validation (Supplemental Fig. 10, 11, 12). To avoid bias due to different experimental protocols and/or datasets, we trained and tested the predictors using external cross validation, i.e. training and testing using different datasets.

We trained the CNN models with 50 filters using the ReLU activation function for hidden layers and linear activation for the output layer. Kernel size was set to 5 and max pooling was set to size of 4. Mean Squared Error (MSE) was used as the loss function and patience for early stopping was set to 10 epochs. Models were regularized by setting the dropout rate to 0.2 to avoid overfitting (Srivastava et al. 2014). We used the R keras package for the implementation of the model (Chollet, Allaire, and Others 2017).

We trained the EN predictor (Friedman, Hastie, and Tibshirani 2010) by setting the value of α to 0.5. The value of the λ was determined using 10-fold internal cross validation. We used the R glmnet package to train the predictor with cv.glmnet function (Friedman, Hastie, and Tibshirani 2009).

Random Forest (RF) predictors were constructed with 500 trees, where each tree was grown with random subsampling of training data using 80% of the training samples. We used the R randomForest package (Liaw and Wiener, n.d.) to build the RF models.

There are different rules for building an ensemble from the individual predictors. The unweighted average approach employed in this study is the mean of the outputs of the individual predictors and does not depend on any prior assumptions on the underlying models. As an alternative, weighted average prioritizes some of the predictors over others based on some prior information. We first calculated the median correlations between predicted gene activity and observed gene expression from the cross validation results and used them as weights for combining the outputs of the predictors. In other words, the predictor that was associated with higher correlation in the training data received higher weight. As another alternative, we used median accuracy (1 - error) as the weight for each predictor.

A more complicated combination rule is the stacked learning (stacking) approach, in which a second layer predictor is trained on the outputs of the first layer predictors (Wolpert 1992; Breiman 1996) using training data. We tested stacking approach with three different second level predictors (belonging to the same family of learners used for the first layer): elastic net regression, artificial neural network and random forest. This approach gave mixed results on the different datasets, offsetting the advantage of stability of ensemble approach. As a result, we decided to employ the unweighted averaging approach, which is the simplest ensemble model and does not rely on any prior information.

### Integration of methylome and transcriptome data using predicted gene activity

We integrated single-cell DNA methylation data with single-cell RNA sequencing data using Seurat version 3 (Stuart et al. 2019). Only cells that have both expression and methylation data were used for the integration. After normalization of both data types, the top 3000 integration features (genes) were selected using the “SelectIntegrationFeatures” function, followed by integration anchors (cells) selected by using the “FindIntegrationAnchors” function, the selected features. The k.filter (number of neighbors) was set to 100 and normalization was set to SCT. The two data types were integrated by using those anchors and the “IntegrateData” function. Finally, we run PCA,UMAP and clustering on the integrated (co-embedded) data including cells from both data types.

### Software availability

The MAPLE software is implemented in R and is freely available under the MIT license. Source code has been deposited at the GitHub repository: https://github.com/yasin-uzun/MAPLE.1.0

## Acknowledgements

We thank the Research Information Services at the Children’s Hospital of Philadelphia for providing computing support. This work was supported by National Institutes of Health of United States of America grants CA226187 and CA233285 (to K.T.), a grant from the Leona M. and Harry B Helmsley Charitable Trust (2008-04062 to K.T.) and a grant from the Alex’s Lemonade Stand Foundation (to K.T.)

## Supplemental Tables

**Supplemental Table 1.** Public single cell sequencing datasets that were used in this study.

| Accession ID | Experimental Protocol | Data modalities (at single cell resolution) | Pubmed ID: Publication |
| --- | --- | --- | --- |
| GSE74535 | scM&T-Seq | Methylome and transcriptome | 26752769: (Angermueller et al. 2016a) |
| GSE121436 | scM&T-Seq | Methylome and transcriptome | 31554804: (Hernando-Herráez et al. 2019) |
| GSE109262 | scNMT-Seq | Methylome, transcriptome and nucleosome occupancy | 29472610: (Clark et al. 2018) |
| GSE121708 | scNMT-Seq | Methylome, transcriptome and nucleosome occupancy | 31827285: (Argelaguet et al. 2019) |
| GSE97179 | snmC-Seq | Methylome | 28798132: Luo et al, Science, 2017 |

## Supplemental Figures

**Supplemental Figure 1.**
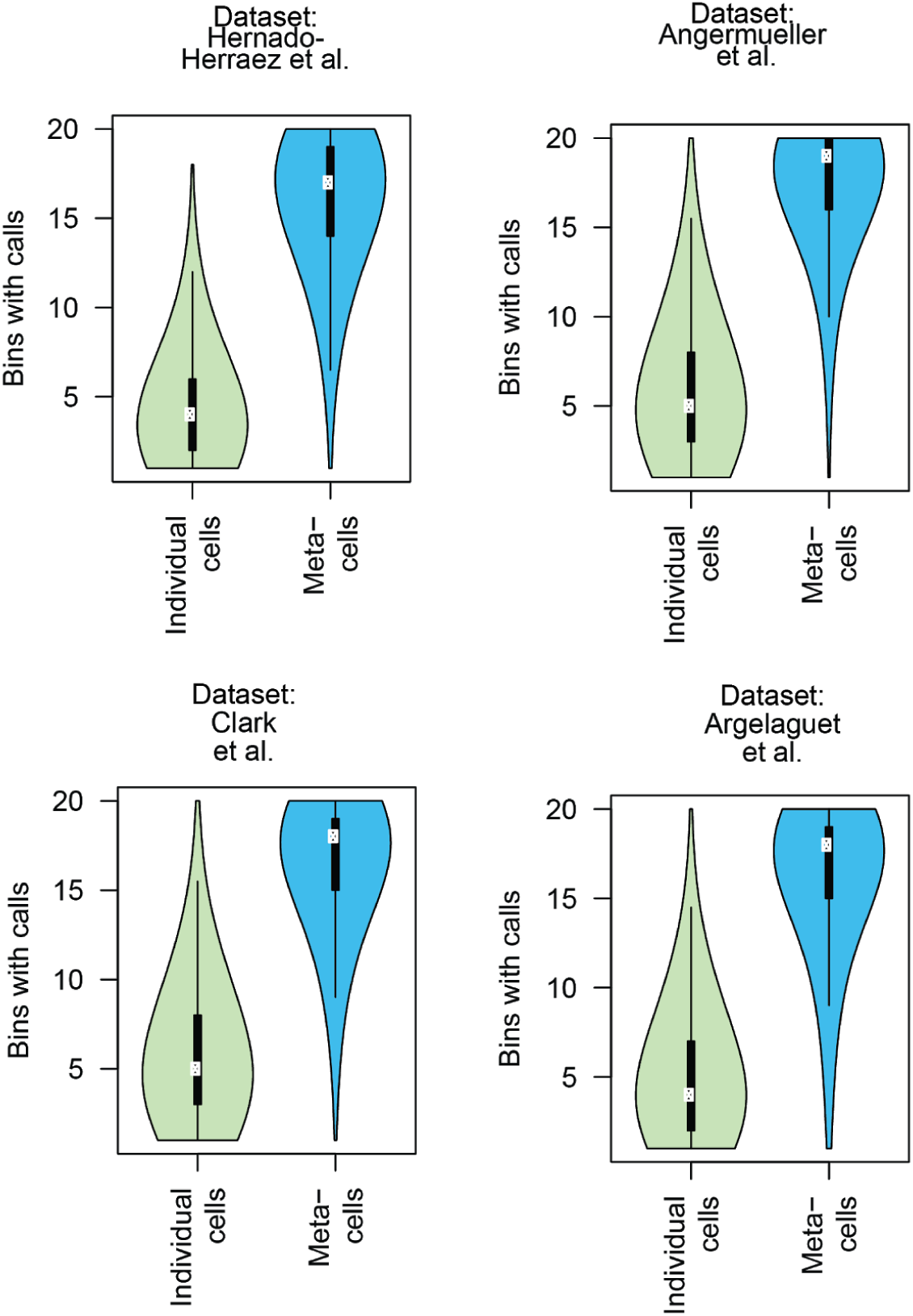
Violin plots showing the effect of using data from meta-cells, instead of individual cells for computation. The +/-5kbp flanking region of TSS was divided into 500bp bins (20 bins). Each data point in plots is a (meta)cell-gene pair and the y-axis shows the number of bins having at least one overlapping cytosine call.

**Supplemental Figure 2.**
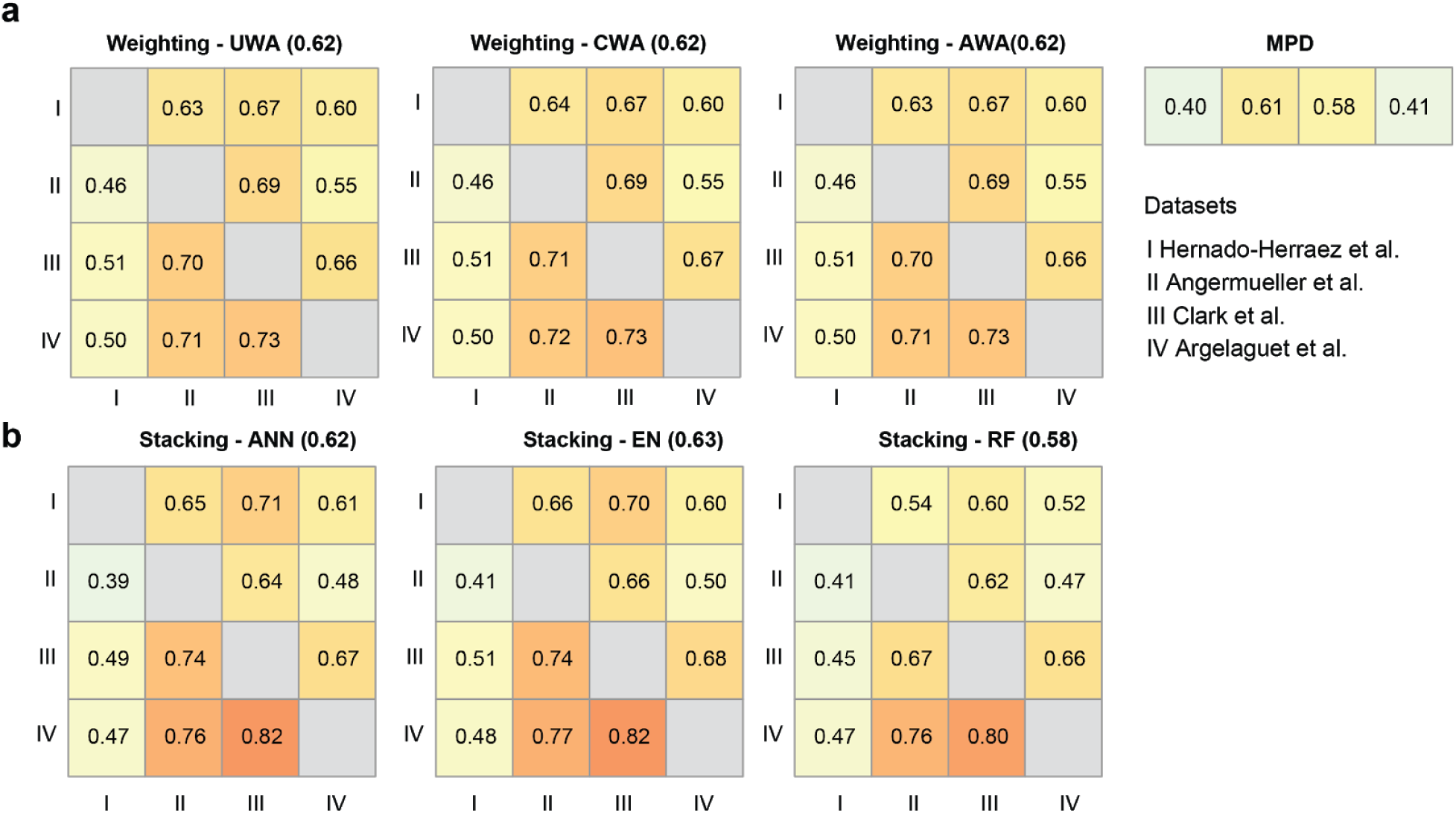
Performance comparison of different ensemble learning combination rules. The rules of combining individual predictors were compared: unweighted (UWA), weighted averaging, and stacking. For weighted averaging, weights were determined either by Spearman correlation coefficient computed with cross-validation (CWA) or by accuracy (1-error) computed with cross-validation (AWA). For stacking, the individual predictors were not given weights. Instead, they were combined into the ensemble by using a second-level predictor. ANN, Artificial Neural Network; EN, Elastic Net Regression; RF, Random Forest. MPD, Mean Promoter De-methylation. **a**) Performance of weighting rules. Heatmap showing global Spearman correlation coefficients between observed gene expression and predicted gene activity for all genes across all cells in a dataset. Rows represent training datasets and columns represent test datasets. **b)** Performance of stacking rules.

**Supplemental Figure 3.**
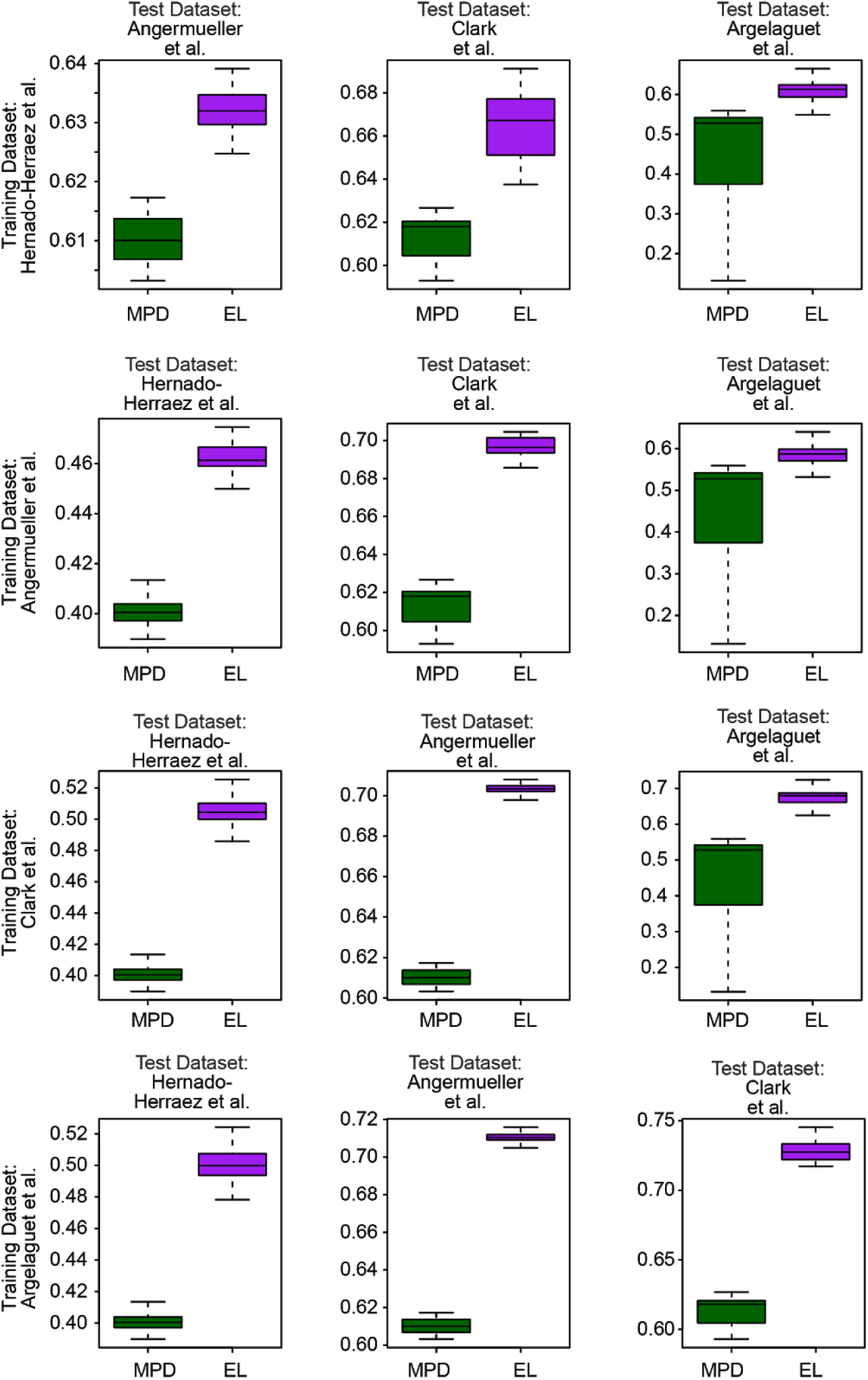
Evaluation of prediction accuracy using Spearman correlation across genes. Each row represents the result using the given training dataset and test datasets. Each boxplot shows the distribution of Spearman correlation coefficients between the predicted gene activity levels and the observed expression levels for the cells in the test dataset. MPD, Mean Promoter De-methylation. EL, Ensemble Learner.

**Supplemental Figure 4.**
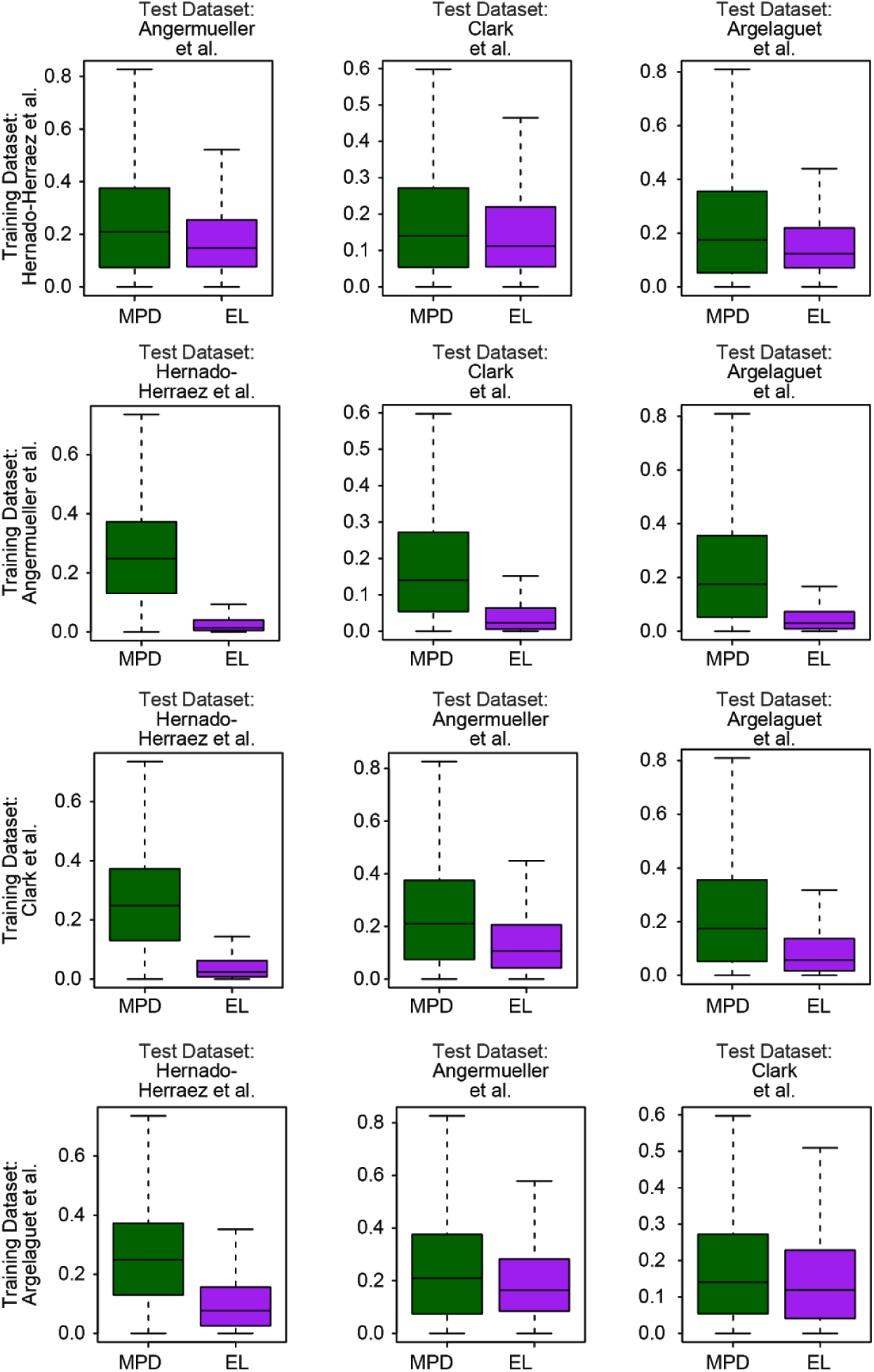
Evaluation of prediction accuracy using median squared error across cells. Each row represents the result using the given training dataset and test datasets. Each boxplot shows the distribution of the median of the squared difference between the predicted gene activity levels and the observed expression levels for each cell. MPD, Mean Promoter De-methylation, EL, Ensemble Learner.

**Supplemental Figure 5.**
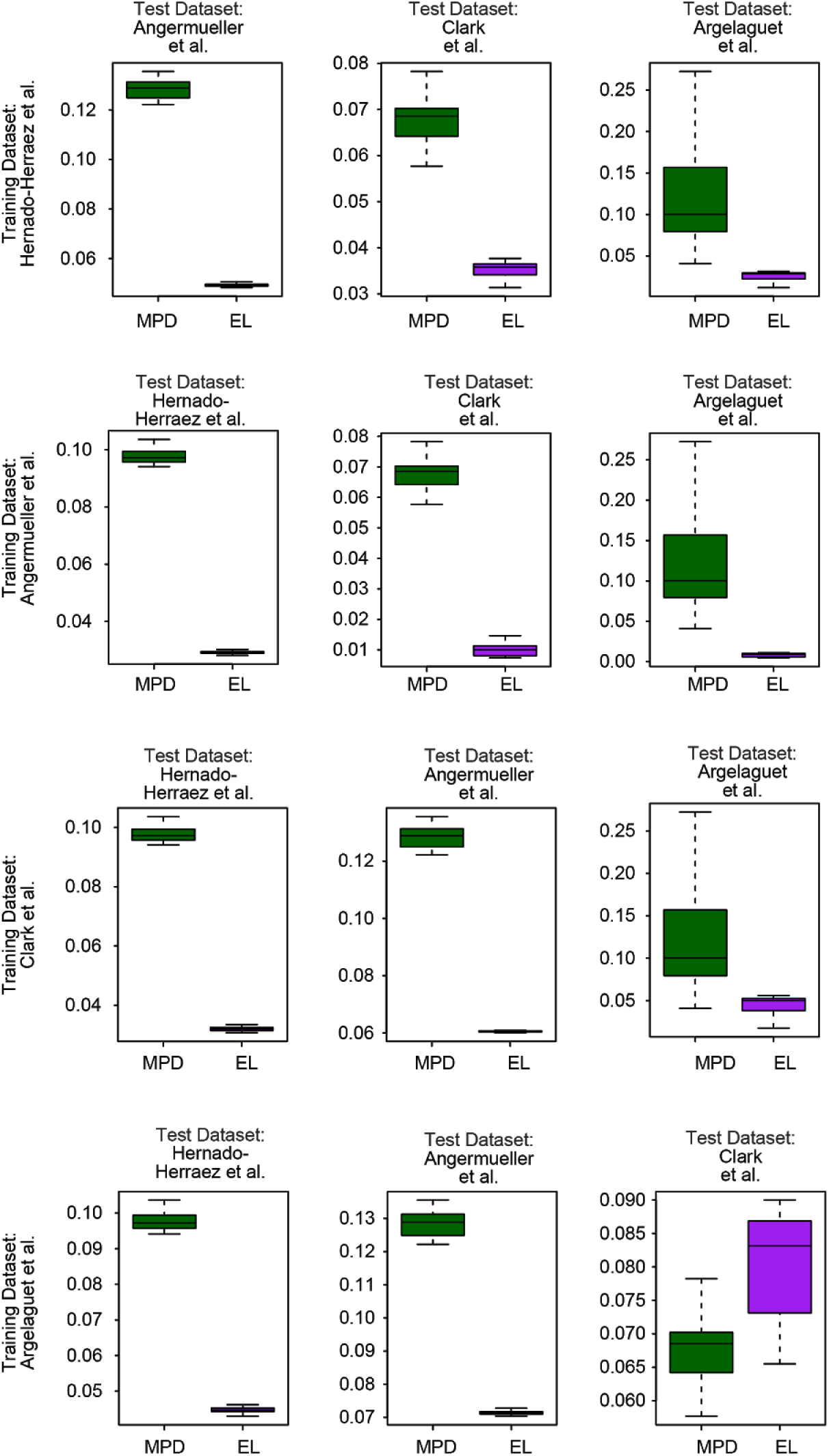
Evaluation of prediction accuracy using median squared error across genes. Each row represents the result using the given training dataset and test datasets. Each boxplot shows the distribution of the median of the squared difference between the predicted gene activity levels and the observed expression levels for each cell. MPD, Mean Promoter De-methylation, EL, Ensemble Learner.

**Supplemental Figure 6.**
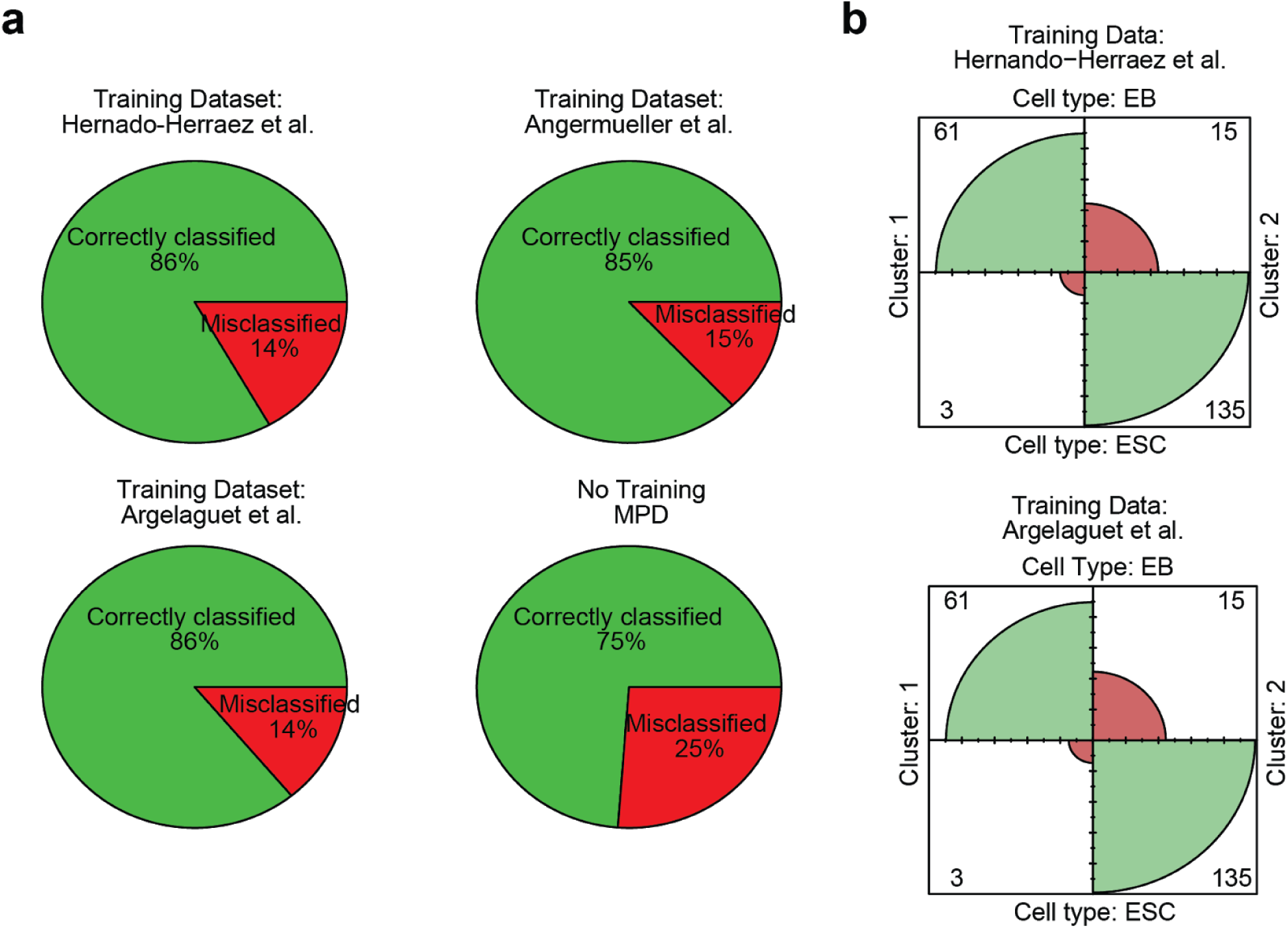
Classification accuracy for cells in the methylome data based on the integration of the dataset of Clark et al. (Clark et al. 2018). Each pie chart shows the percentage of cells in DNA methylation data that were assigned to the correct cell type (EB or ESC) using k-Nearest Neighbor (kNN) classification using nearest cells from scRNA data in the integrated PCA. Bottom right chart shows the percentage of correctly classified cells when mean promoter de-methylation was used for the integration. Remaining three panels show the percentage of correct classification with ensemble method using three different training sets. Supervised predictor outperformed the unsupervised mean de-methylation based predictor, regardless of the training dataset. **b)** Confusion matrices for the class assignments of EB and ES cells. Cluster 1 is composed of EBs and Cluster 2 is composed of ESCs. Ensemble method was used for constructing a gene activity matrix from DNA methylation data and then scRNA and gene activity matrices were integrated with Seurat (Stuart et al. 2019). Each confusion matrix is the result of integration using the gene activity matrix predicted by the models trained by the datasets shown at the top.

**Supplemental Figure 7.**
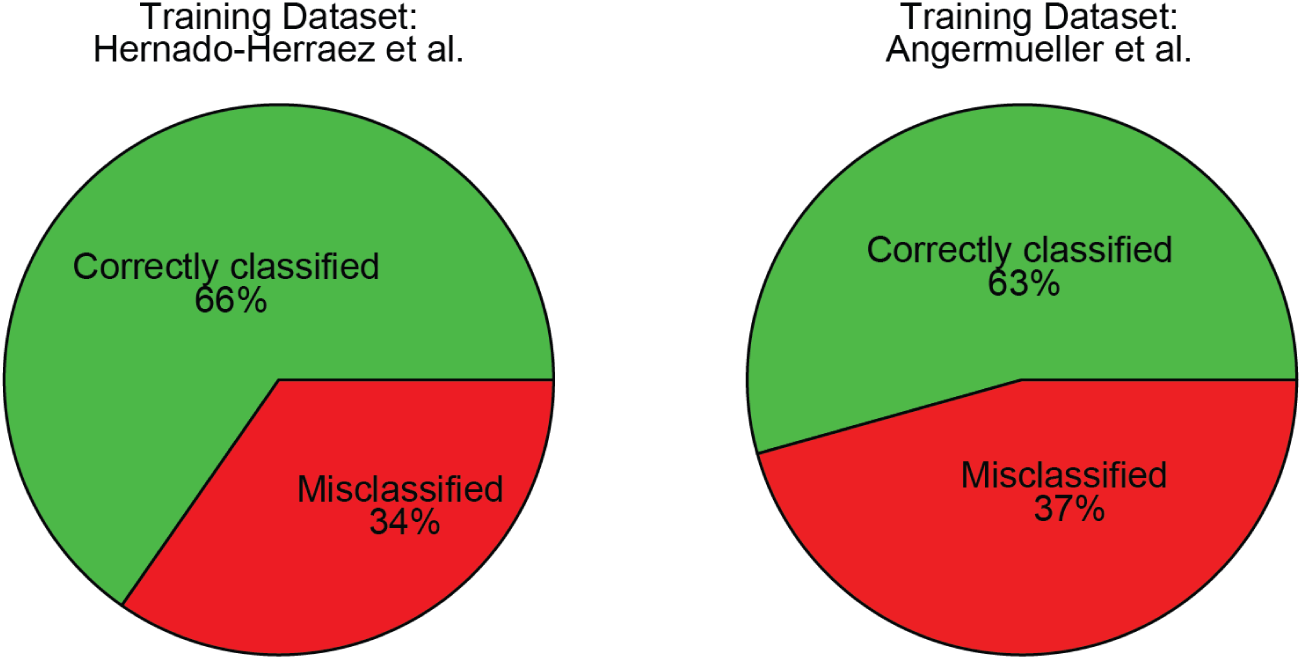
Percentage of correct classification for the cells in the methylome data based on the integration on the dataset of Argelaguet et al (Argelaguet et al. 2019). Each pie chart shows the percentage of cells in DNA methylation data that were assigned to the correct cell type (same embryonic day) using k-Nearest Neighbor (kNN) classification using cells from scRNA data in the integrated PCA. The models were trained with the dataset shown on top.

**Supplemental Figure 8.**
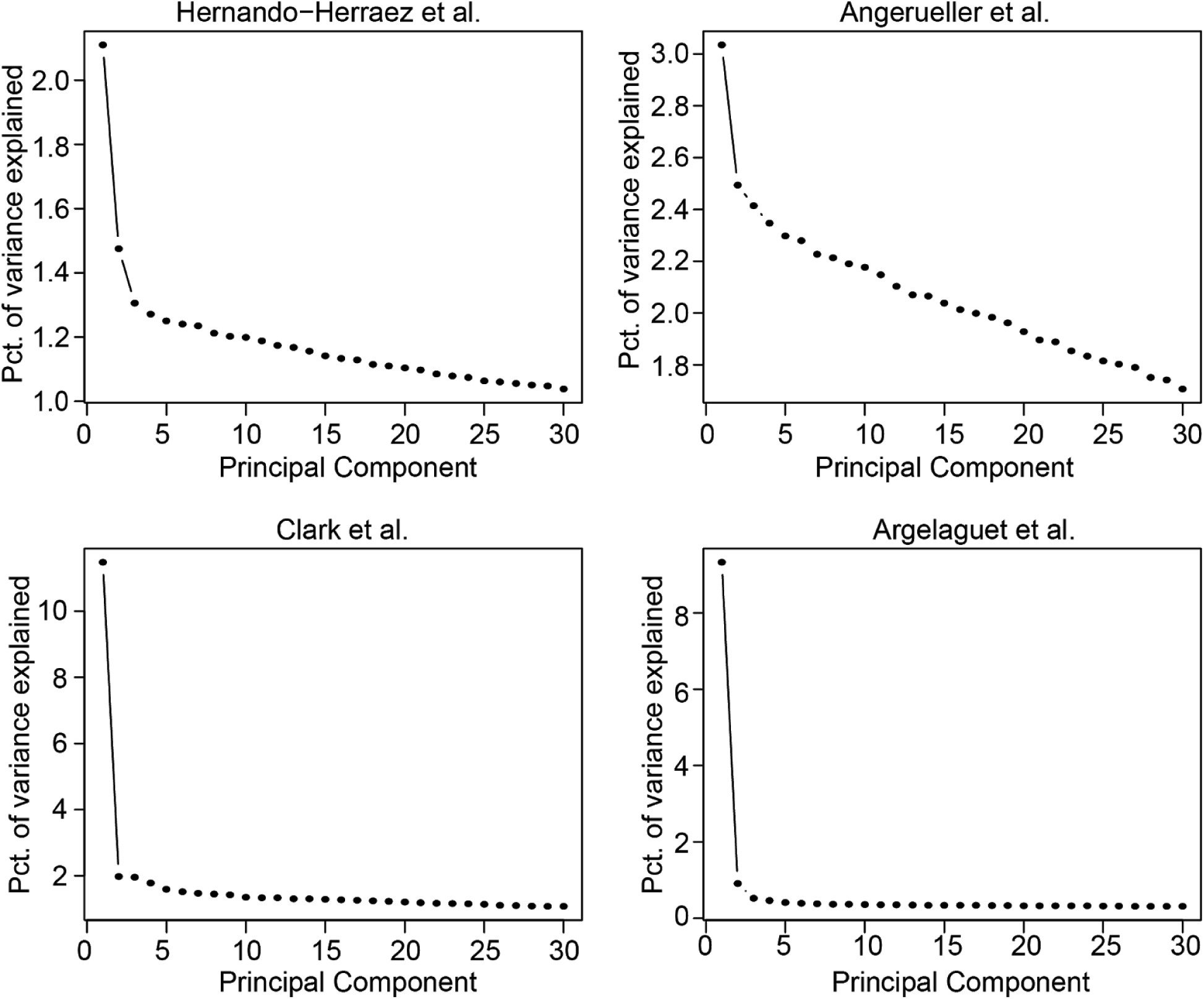
Elbow plots showing the percentage of variance in the data explained by each principal component in the principal component analysis, sorted from highest to lowest.

**Supplemental Figure 9.**
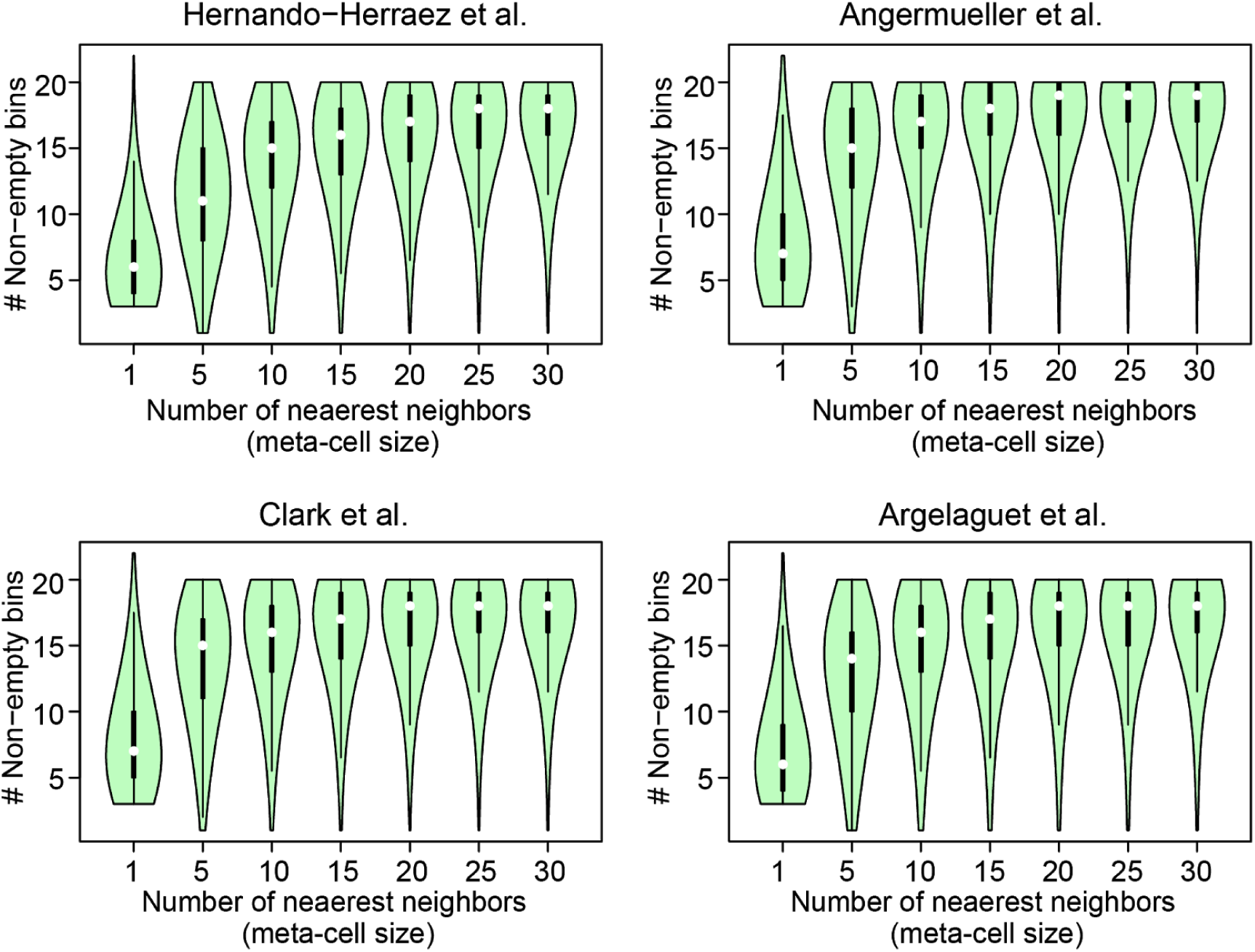
Violin plots showing the distribution of the number of non-empty bins for individual cells (*k*=1) and meta-cells of varying sizes. Each data point is one cell (k=1) or meta-cell (k>1), and the value is the mean number of non-empty bins for the promoters of that cell. White circle shows the median value. Black rectangle shows the upper and lower quartiles. Whisker shows 1.5 times the interquartile range.

**Supplemental Figure 10.**
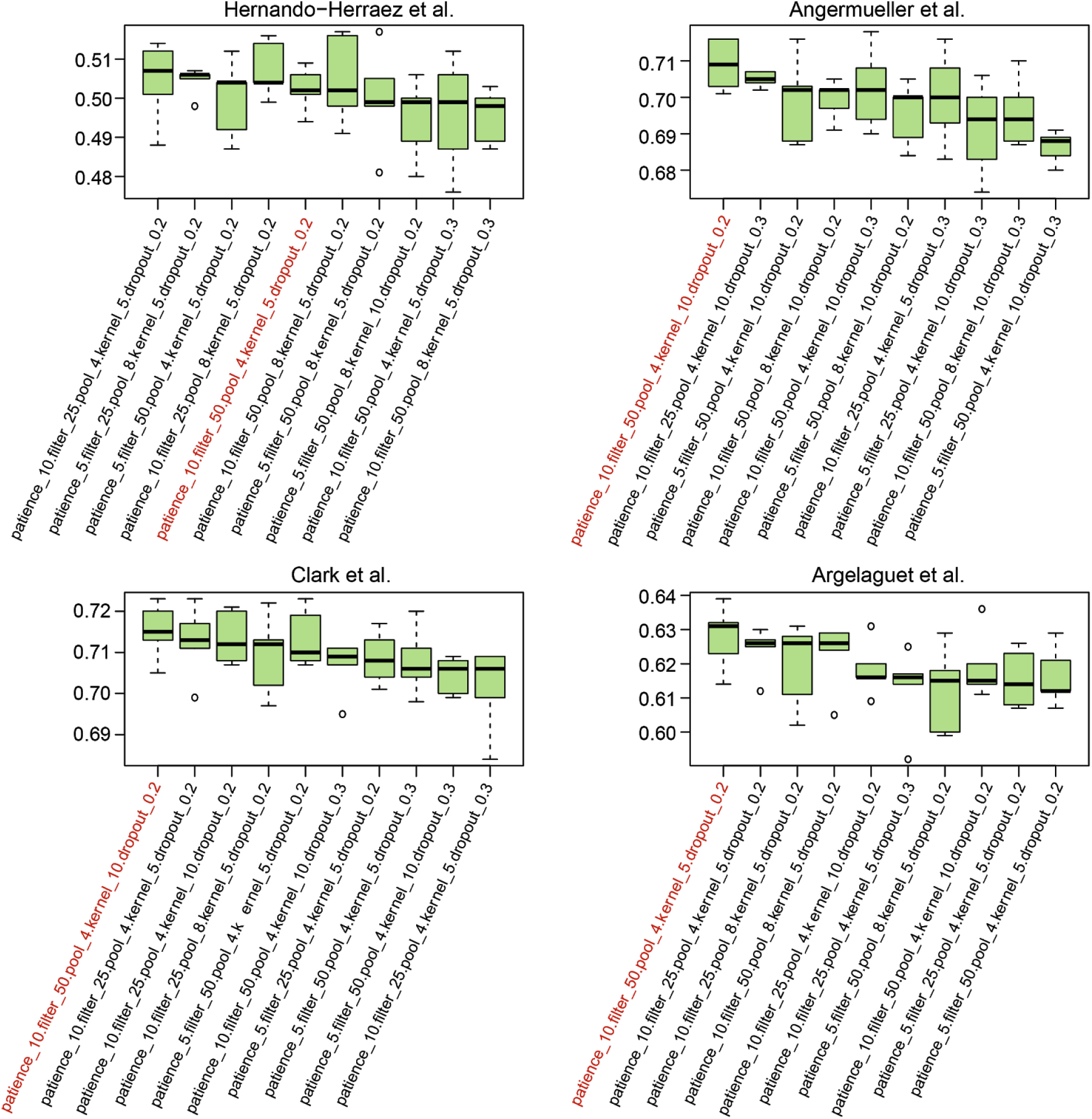
Box plots showing the parameter tuning results with 5-fold cross validation for convolutional neural network predictor. Top 10 parameter settings with highest Spearman correlation (median of 5 runs) between the predicted gene activity and observed expression for all the cells and genes are shown. Each data point is a single run. Parameter name is followed by the value for each setting. Patience, number of epochs to stop if there is no reduction in error during optimization step; filter, number of filters; pool, pooling size; kernel, kernel size; dropout, dropout rate. Parameter setting used in the ensemble predictor is highlighted in red.

**Supplemental Figure 11.**
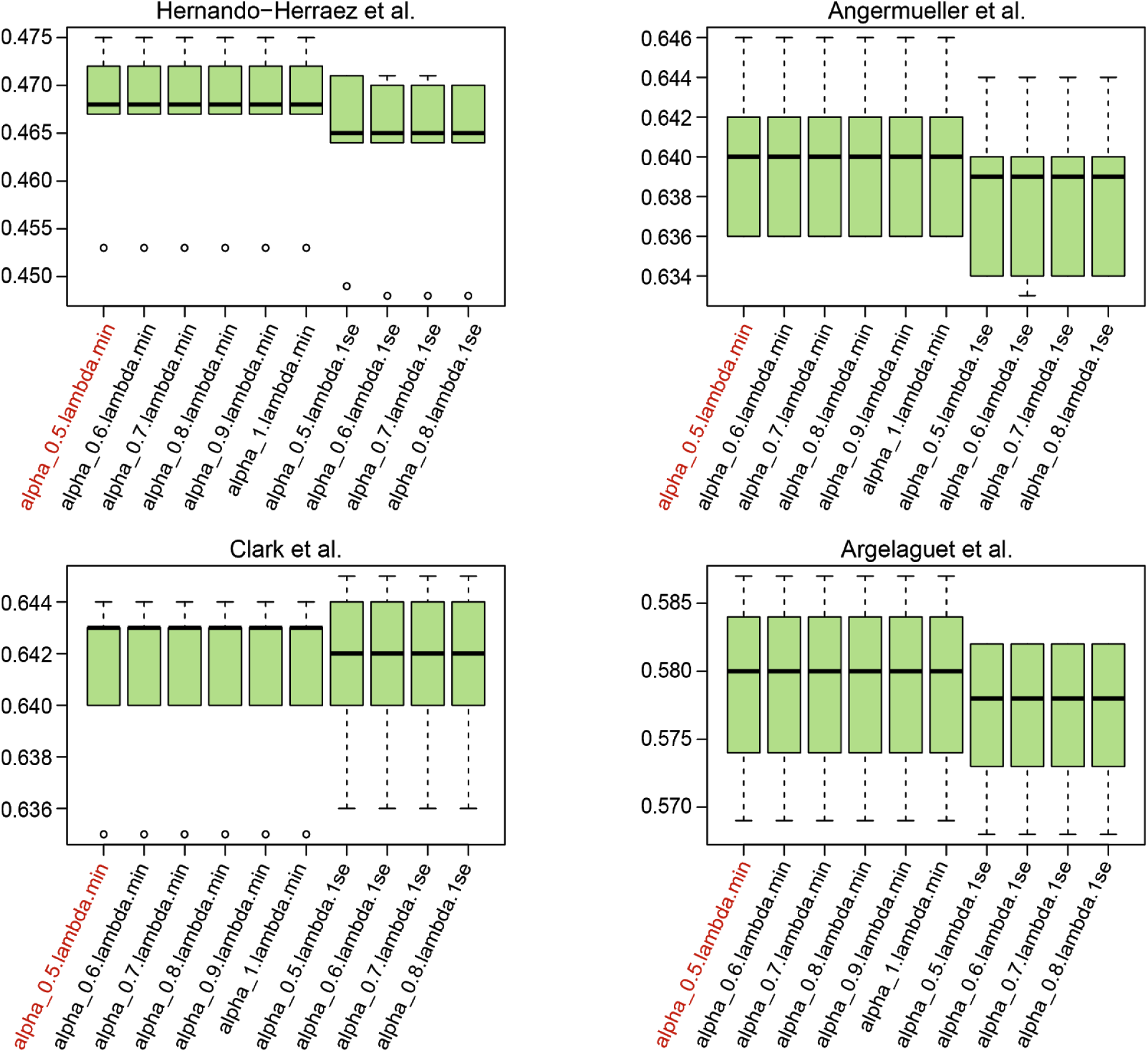
Box plots showing the parameter tuning results with 5-folds cross validation for elastic net predictor. Top 10 parameter settings with highest correlation are shown. Each data point is a single run. Parameter name is followed by the value for each setting. alpha, alpha value setting the balance between LASSO and Ridge regressions. lambda.min, choice of lambda that gives the minimum error via internal cross validation. lambda-1se, choice of lambda in one standard error vicinity of the lambda.min, for regularization. Parameter setting used in the ensemble predictor is highlighted in red.

**Supplemental Figure 12.**
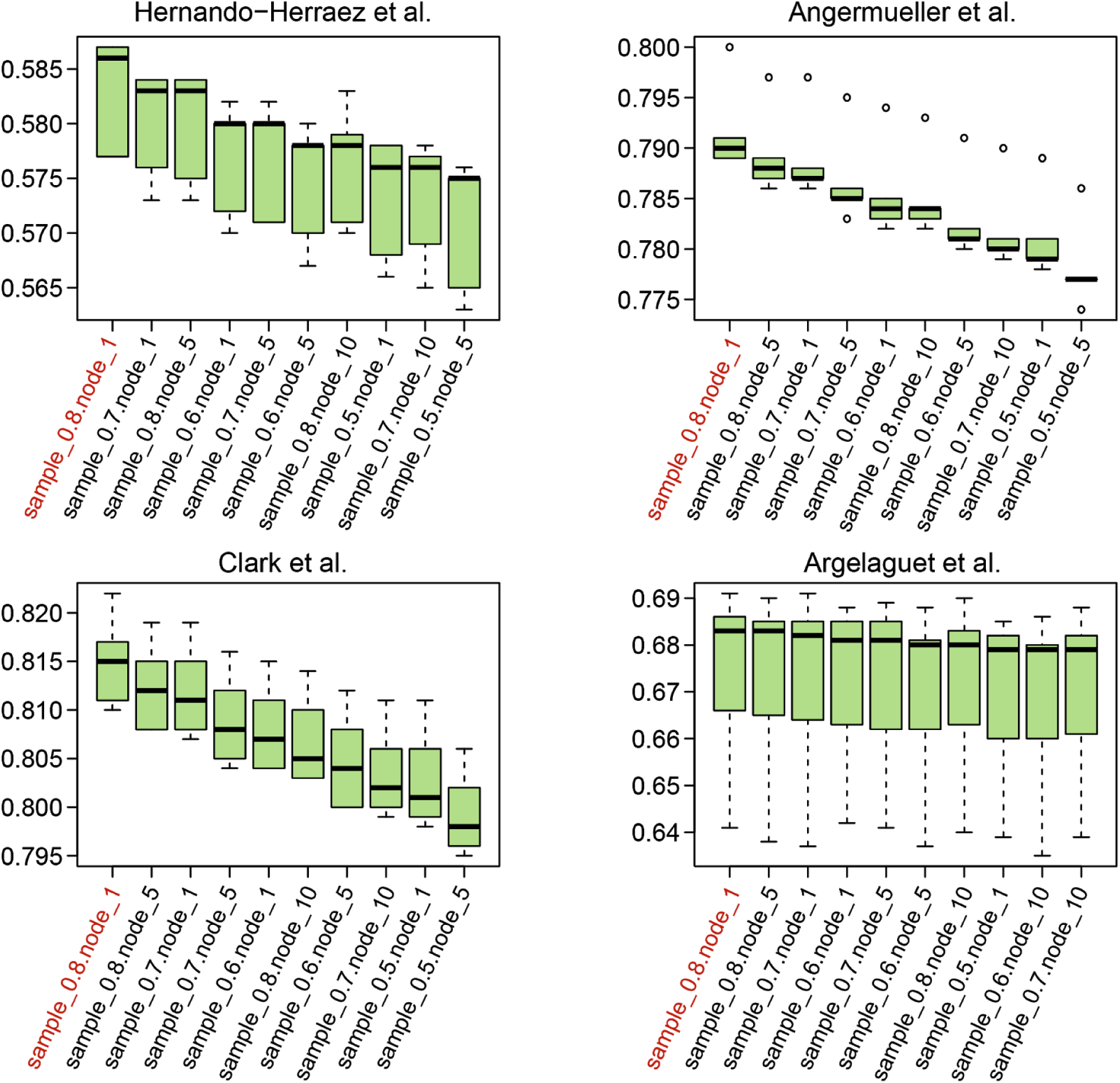
Box plots showing the parameter tuning results with 5-folds cross validation for random forest predictor. Top 10 parameter settings with highest correlation are shown. Each data point is a single run. Parameter name is followed by the value for each setting. sample, ratio of the input samples to be used for constructing each tree. node, number of minimum terminal nodes. Parameter setting used in the ensemble predictor is highlighted in red.

